# Loss of 18q alters TGFβ signalling affecting anteroposterior neuroectodermal fate in human embryonic stem cells

**DOI:** 10.1101/2024.10.09.617397

**Authors:** Yingnan Lei, Mai Chi Duong, Nuša Krivec, Charlotte Janssens, Marius Regin, Anfien Huyghebaert, Edouard Couvreu de Deckersberg, Karen Sermon, Diana Al Delbany, Claudia Spits

## Abstract

Chromosomal abnormalities acquired during cell culture can compromise the differentiation potential of human pluripotent stem cells (hPSCs). In this work, we identified a diminished differentiation capacity to retinal progenitor cells in human embryonic stem cells (hESCs) with loss of chromosome 18q. Time-course gene-expression analysis during spontaneous differentiation and single-cell RNA sequencing found that these variant cell lines poorly specified into anterior neuroectoderm, and, when progressing through differentiation, they yielded poorly pigmented cells, with proliferating and pluripotent cell populations. The variant cell lines showed dysregulation of TGFβ signaling during differentiation, and chemical modulation of the TGFβ pathways showed that the basis of the improper specification was due to imbalances in the anteroposterior neuroectodermal fate commitment.

## Introduction

The Retinal Pigment Epithelium (RPE) is a monolayer of post-mitotic, cobblestone-shaped, pigmented, and polarized cells positioned at the interface between the choriocapillaris and the sensory neural retina. The differentiation of this tissue begins shortly after gastrulation, first with the specification of the eye field from the anterior neural plate. After, the optic vesicles appear, and the RPE will emerge from their proximal part. Dysfunction, degeneration, and loss of RPE cells are prominent features of retinal degenerative disease, such as Stargardt disease (SD) and age-related macular degeneration (AMD). These conditions frequently result in significant vision loss and ultimately result in blindness ^1,2^. Phase I/II clinical transplantation trials utilizing human pluripotent stem cell-derived RPE (hPSC-RPE) for the treatment of AMD and SD have robustly demonstrated both safety and feasibility ^3–8^.

Human pluripotent stem cells (hPSCs) are prone to acquire genomic abnormalities, which may undermine their suitability for clinical applications ^9^. The most common recurrent mutations observed in hPSCs include gains of chromosomes 1q, 12p, 17, 20, X, and to a lesser extent, losses of chromosomes 10p, 18q, and 22p, with point mutations in *TP53* also frequently appearing ^9–14^. A critical concern regarding these genetic variants is whether and how they alter the differentiation capacity of hPSCs, thereby potentially priming differentiated cells for malignant transformation ^9,15–18^.

In previous work, we found that loss of 18q impairs directed neuroectodermal differentiation in human embryonic stem cells (hESCs^19^). In this study, we aimed to elucidate the impact of a loss of 18q on the developmental trajectory that leads to the formation of the RPE, with a special focus on the critical step of specifying the anterior and posterior plate of the neuroectodermal lineage. Further, we dissected characteristics of the derived precursors and mature RPE. To achieve this, we compared the differentiation capacity towards RPE of hESC lines with 18q deletions (hESCs^del18q^) to genetically balanced lines (hESCs^WT^). Single-cell RNA sequencing (scRNA-seq) was employed to characterize cellular diversity and assess variations between hESCs^WT^ and hESCs^del18q^. Finally, treatment with specific inhibitors of key developmental pathways was used to unravel the mechanisms influencing the differentiation process.

## Materials and methods

### hESCs maintenance and passaging

All s lines were derived and characterized as reported previously ^20,21^ and are also registered in the EU hPSC registry (https://hpscreg.eu/). hESCs were maintained in NutriStem hESC XF medium (NS medium; Biological Industries) with 100 U/mL penicillin/streptomycin (P/S) (Thermo Fisher Scientific) in a 37 °C incubator with 5% CO_2,_ and the culture medium was changed daily. The tissue culture dishes and plates (Thermo Scientific) were coated with 10 µg/mL Biolaminin 521 (Biolamina®) at 4°C and then incubated at 37 °C for at least 30 min before the cells were seeded. The medium was supplemented with 10 μM Rho kinase (ROCK) inhibitor Y-27632 (ROCKi, Tocris) for the first 24 h after passaging.

### hESCs-RPE differentiation

The induction of RPE differentiation was initiated using a slightly modified version of the protocol from a previous publication ^22–24^. hESCs were plated at 100,000 cells/cm^2^ on 20 μg/ml laminin-521 coated dishes or plates with 10 μM ROCK inhibitor Y-27632 during the first 24 h in NutriStem hESC XF medium. When reaching to 90% confluence, the medium was replaced with NutriStem® hPSC XF GF-free medium without basic fibroblast growth factor (bFGF) and transforming growth factor (TGFβ) with media changed every day until the timepoint for sample collection without re-plaiting during the whole differentiation process. The pigmentation started to be visible from week 4 and pigmented areas were mechanically cut out using a sharpened glass pipette at around 90 days. The replated cells were fed twice a week with NutriStem® hPSC XF GF-free medium without bFGF and TGFβ.

### Copy number variant (CNV) analysis

The genetic content of the hESCs was assessed through shallow whole-genome sequencing by the BRIGHTcore of UZ Brussels, Belgium, as previously described ^25^.

### RNA isolation and cDNA synthesis

RNA was isolated using RNeasy Mini and Micro kits (Qiagen) following the manufacturer’s guidelines, including on-column DNase I treatment. A minimum of 500 ng of mRNA was reverse-transcribed into biotinylated cDNA using the First-Strand cDNA Synthesis Kit (Cytiva) with the NotI-d(T)18 primer.

### Quantitative real-time PCR (qRT-PCR) for gene expression analysis

Quantitative real-time PCR (qRT-PCR) was carried out using TaqMan mRNA expression assays (Thermo Fisher Scientific) and TaqMan 2× Mastermix Plus – Low ROX (Eurogentec) on a ViiA 7 thermocycler (Thermo Fisher Scientific) using the standard cycling protocol provided by the manufacturer. The relative expression of target genes was quantified using the comparative threshold cycle (Ct) method and normalized to the TaqMan *GUSB* transcript (Applied Biosystems) as the endogenous housekeeping gene. All the samples were run in triplicate, and the related TaqMan assays used in the present study are listed in Supplementary Table 2.

### Immunostaining

Differentiated cells were first fixed in a solution of PBS containing 3.7% formaldehyde (Sigma-Aldrich) for 15 min, permeabilized in 0.1% Triton X for 10 min (Sigma-Aldrich) and then blocked with 10% fetal bovine serum (ThermoFisher Scientific) for 1 h at room temperature (RT). Sequentially, primary antibodies appropriately diluted in blocking solution (1:200 dilution in 10% FBS) were incubated overnight at 4 °C. Thereafter, secondary antibodies conjugated to Alexa 488, Alexa 594 (1:200 dilutions in 10% FBS, Thermo Fisher Scientific) and Hoechst (12000 dilution, ThermoFisher Scientific) were applied for 1-2 h at room temperature in the dark. Confocal images were acquired under an LSM800 (Carl Zeiss) confocal microscope at 20x magnification. The lists with antibodies can be found in supplementary Table 3.

### Single-cell suspension preparation

Following 2 months of RPE differentiation, the RPE cells were washed at least 3 times with PBS to remove the dead cells and dissociated using Papain (Worthington Biochemical Corporation) for 30mins-1h at 37 °C following the manufacturer’s guidelines. After dissociation, the cell concentration and viability was measured using Annexin V & Propidium Iodide (Thermo Fisher Scientific). Aprroximately 4 million cells were collected and fixed according to the manufacturer’s protocol (Parse Biosciences). After fixation, the cell concentration was measured again for each suspension before being stored at -80 °C. Cells were then subjected to the Single Cell Whole Transcriptome Kit v2 (Parse Biosciences) for library construction. Barcoding and sequencing library generation were performed according to the manufacturer’s protocol and libraries sequenced using the high-throughput NovaSeq (Illumina) with 20k reads per cell.

### Sc-RNA sequencing

Parse data was aligned with the ParseBiosciences-Pipeline v0.9.6 p using the GRCh38 reference (refdata-gex-GRCh38-2020-A). Further analysis of the raw count matrices was loaded into the R package Seurat (5.3.0) for downstream analyses.

Dimension reduction was performed using RunPCA() and RunUMAP functions. The top principal components (PCs) were used to construct nearest-neighbor graphs and identify cell clusters using the FindNeighbors() and FindClusters() functions of the Seurat R package. Single-cell data were visualized by the “Dimplot” function. “VlnPlot” functions were used to display the RPE related gene expression levels. “DoHeatmap” functions were applied to show the heatmap. “AddModuleScore_UCell” and “DotPlot” functions were employed for the pigmentation genes expression.

### GSEA analysis

Highly Differentially expressed genes of each cell cluster were analyzed using the Seurat “FindAllMarkers” function using “MAST” test method and genes were ranked by fold change in expression level. GSEA process analysis was performed via the clusterProfiler R package and GSEA function and the Molecular Signature Database C2 and H (MSigDB).

### SCENIC Analysis

SCENIC analysis was carried out following the SCENIC command line protocol (Aibar et al., 2017). AUCell used the area under the curve to calculate the enrichment of the regulon across the ranking of all genes, resulting in a matrix of the activity of each regulon in each cell cluster. Downstream analyses were done in R combing the cell cluster information obtained from Seurat. Regulon specificity scores (RSS) were computed based on the cell populations clusters identified by Seurat and the AUC heatmap was plotted by the ComplexHeatmap::Heatmap function.

### Statistical analysis

All differentiation experiments were carried out in at least triplicate (n ≥ 3). All data are presented as the mean ± standard error of the mean (SEM). Statistical evaluation of differences between 2 groups was performed using unpaired two-tailed t tests in GraphPad Prism 9 software, with p < 0.05 determined to indicate significance.

## RESULTS

### hESCs^del18q^ exhibit impaired anterior neuroectoderm induction and poor progression through poor eye field induction and RPE specification

To assess the effects of the 18q deletion on RPE differentiation, two hESC lines carrying a loss of chromosomes 18 (hESCs^del18q^: VUB13^del18q^ and VUB14^del18q^) and three genetically balanced lines (hESCs^WT^: VUB02^WT^, VUB03^WT^ and VUB14^WT^) were differentiated into RPE ^24^. The karyotypes were confirmed through shallow whole genome sequencing, and the cell lines were utilized within 10 passages post-thawing to minimize the risk of (additional) genetic changes. Additionally, DNA samples were collected at the initiation of RPE differentiation to assess that the lines did not acquire the highly recurrent gains of 1q, 12p, 20q11.21, and 17q, by quantitative real-time PCR.

Following three months of RPE differentiation induction, extensive regions of the culture dishes in WT cell lines were populated with pigmented cells, a hallmark of successful differentiation to RPE. By contrast, hESCs^del18q^ exhibited only isolated and sparse patches of pigmentation, indicating a substantially diminished RPE differentiation capacity (Fig. 1A). The lower differentiation efficiency was also confirmed by mRNA expression analysis of neuroectodermal (*PAX6*), RPE (*PMEL*, *MITF*, *RPE65*, *BEST1*), and pluripotency (*NANOG* and *POU5F1*) markers (Fig. 1B and Sup. Fig. 1A). All the RPE markers showed a strong induction in hESCs^WT^, whereas hESCs^del18q^ exhibited significantly lower expression levels, consistent with the observed colony pigmentation patterns on the culture dishes. VUB14^del18q^ still retained expression of pluripotency markers after three months of differentiation (Fig. 1B and Sup. Fig. 1A).

**Figure 1.**
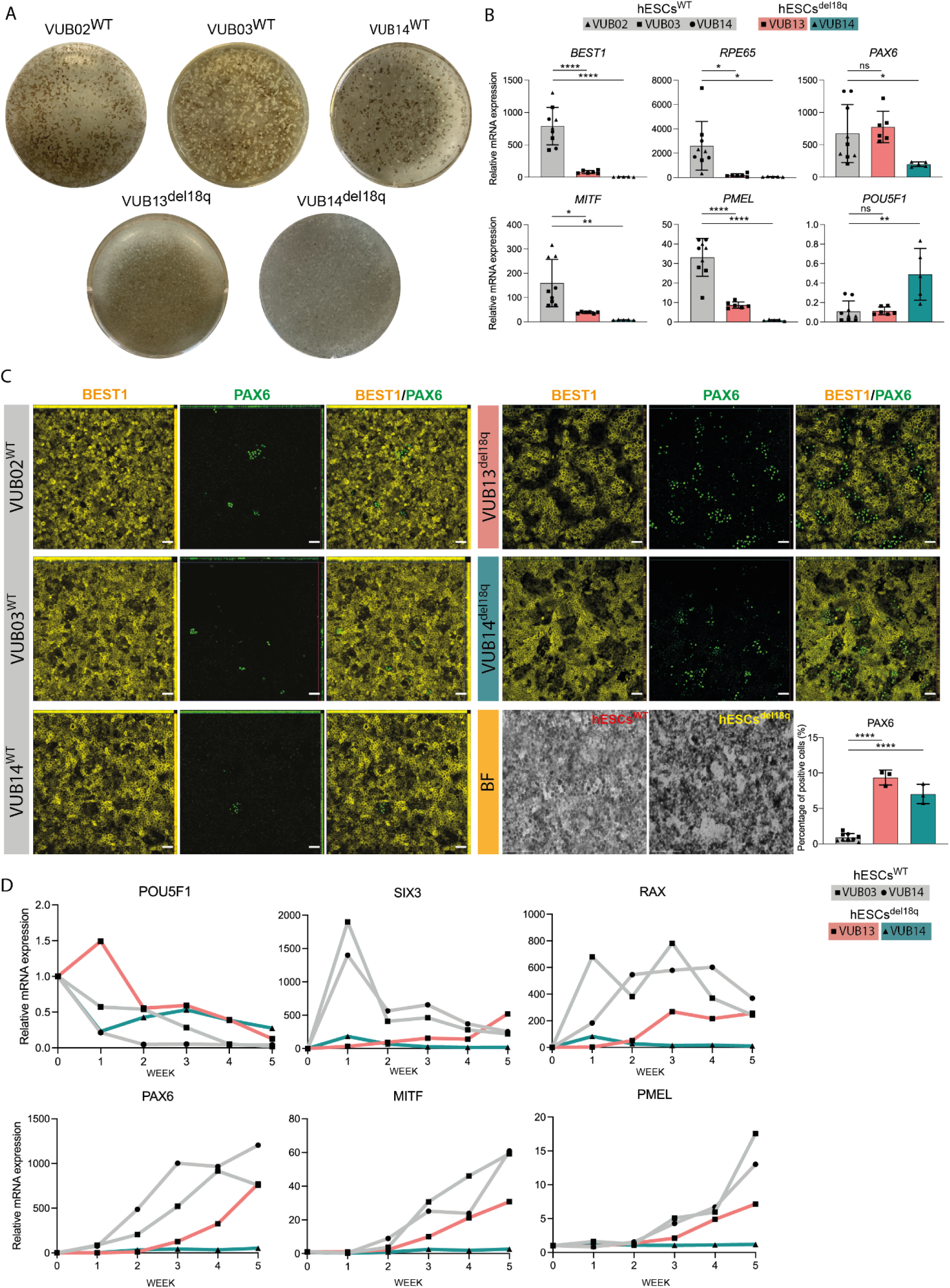
hESCs with a loss of 18q show a reduced differentiation potential to RPE and an overall poor anterior neuroectoderm induction. **A.** Whole culture well images to illustrate the pigmentation observed after 3 months-differentiation of hESCs^WT^ and hESCs^del18q^. B. Relative mRNA expression of neuroectodermal/retinal progenitor (*PAX6*), RPE (*RPE65*, *BEST1*, *MITF*, *PMEL*) and pluripotency (*POU5F1*) markers of the dishes shown in A, prior to RPE purification. Data information: Data are shown as the means ± SEM (hESCs^WT^: N=3/per line, VUB13^del18q^: N = 6, VUB14^del18q^: N = 5). C. Staining of purified RPE for the mature RPE marker BEST1 and the neuroectodermal marker PAX6, scale bars of the are 50 µm. BF (bright field) images together with counting of PAX6 positive cells (hESCs^WT^: N=3/per line, VUB13^del18q^: N = 3, VUB14^del18q^: N = 3). D. Temporal dynamics trajectory of spontaneous RPE differentiation, with relative mRNA expression of early RPE (*MITF*, *PMEL*), retinal progenitor (*PAX6, RAX* and *S I X*)*3*and pluripotent state (*POU5F1*) markers at 5 time points (W1, W2, W3, W4, W5). Each datapoint refers to an independent differentiation experiment and *, **, *** and **** represent statistical significance between samples at 5%, 1%, 0.1% and 0.01% respectively (unpaired t-test).

In order to further characterize the differentiated cells, the pigmented foci obtained after 90 days of differentiation were manually dissected and replated to produce a pure population of RPE cells (brightfield image in Fig.C). Immunostaining for BEST1, ZO-1, PMEL and PAX6 revealed that the purified RPE population obtained from hESCs^del18q^ carries a higher proportion of PAX6-positive precursor cells and a diminished population of mature RPE cells. In hESCs^WT^-derived cells, we observed an average of 0.9±0.4 % of PAX6 positive cells, as compared to 8.1±1.6% in hESCs^del18q^ (Fig. 1C). The same pattern was found in the ZO-1 and PMEL staining, which displayed negative areas in the hESCs^del18q^-derived RPE, contrasting with the homogeneously positive staining in the hESCs^WT^-derived RPE (Sup. Fig. 1B). This suggests that the mutant lines are unable to generate a fully mature RPE.

Subsequently, we aimed at understanding if the lower degree of maturity of the cells was due to the intrinsic incapability to generate mature RPE or if there was a delay in the specification of the early eye field progenitors. We performed a time-course experiment, collecting weekly samples from differentiating cells over a period of 5 weeks, and analyzed the expression of pluripotency-associated genes (*NANOG*, *POU5F1*), early eye field progenitor genes (*RAX*, *SIX3*), a pan-neuroectoderm marker (*PAX6)*, early RPE (*MITF*, *PMEL*) and late RPE (*BEST1*, *RPE65*) markers (Fig. 1D and Sup. Fig. 1C). In hESCs^WT^, we observed a progressive downregulation of the pluripotency-associated genes with a concomitant increase of the progenitor markers, culminating in the induction of expression of RPE markers in the final two weeks of the experiment. The hESCs^del18q^ lines showed a different pattern of expression. VUB13^del18q^ exhibited an overall delay in the repression of pluripotency markers, and induction of lineage-specific genes, with lower levels of expression of RPE genes at the end point. VUB14^del18q^ never fully repressed the pluripotency genes and presented very poor induction of progenitor/RPE markers. It is worth noting that, despite the line-specific profiles, both hESCs^del18q^ lines had significantly low induction of the anterior neuroectoderm marker *SIX3* and of the early eye field gene *RAX*. Taken together, this shows that hESCs^del18q^ do not only fail to efficiently differentiate into anterior neuroectoderm, required for eye field formation, but that if they do progress to that stage, they have a reduced capacity for RPE commitment.

### hESCs^del18q^ differentiate into cell populations containing residual undifferentiated cells, proliferating cells and immature RPEs

To gain deeper understanding on this differentiation impairment, we characterized the cells obtained after 60 days of spontaneous differentiation from hESCs^WT^ and hESCs^del18q^ by single cell RNA sequencing (scRNA-seq). The cell clusters were annotated based on cluster-specific enriched markers that aligned with published cell lineage markers (listed in Sup. Table1).

Clustering analysis was performed independently for each sample, revealing five populations in common between the two hESCs^WT^-derived cells: RPE cells, retinal progenitor cells (RPC), Amacrine Cells or retinal ganglion cells (AC/RGC), Cortical Hem (CH) and Amnion cells (Fig. 2A). The expression of the markers for each cluster is shown in Fig. 2B. The samples from the hESCs^del18q^-derived cells overall showed similar cell populations as hESCs^WT^, but with different proportions (Fig. 2A). The largest cluster of VUB13^del18q^ was marked by high expression levels of PDGFRA, SFRP2, CP, PCDH19, and VAV3. This cluster is unique to this subline, and is likely composed of Müller glia cells. Notably, 48% and 46% of cells were categorized as RPE in the WT cells, compared to 29% and 36% in the two mutant lines (Fig. 2A), suggesting a reduced RPE cell population in the two hESCs^del18q^. Conversely, the levels of RPE-related genes were significantly lower in the RPE subpopulations of hESCs^del18q^ as compared to hESCs^WT^ (Fig. 2C), and particularly, the pigmentation-related genes, in line with the poor pigmentation visually observed in the hESCs^del18q^-derived RPE (Fig.2D, pigmentation related genes listed in Sup. Table1). hESCs^del18q^ also exhibited higher proportions of proliferating cells compared to hESCs^WT^, with proliferating cells accounting for 12% and 25% in hESCs^del18q^, versus only 4% in VUB03^WT^ and none in VUB02^WT^ (Fig. 2A). Moreover, VUB14^del18q^ exhibited a cluster that remained in the pluripotent stage, in line with the retention of pluripotency-associated gene expression found at the end point of the time-course experiment shown in figure 1D and the 3 months RPE induction in Figure 1B.

**Figure 2.**
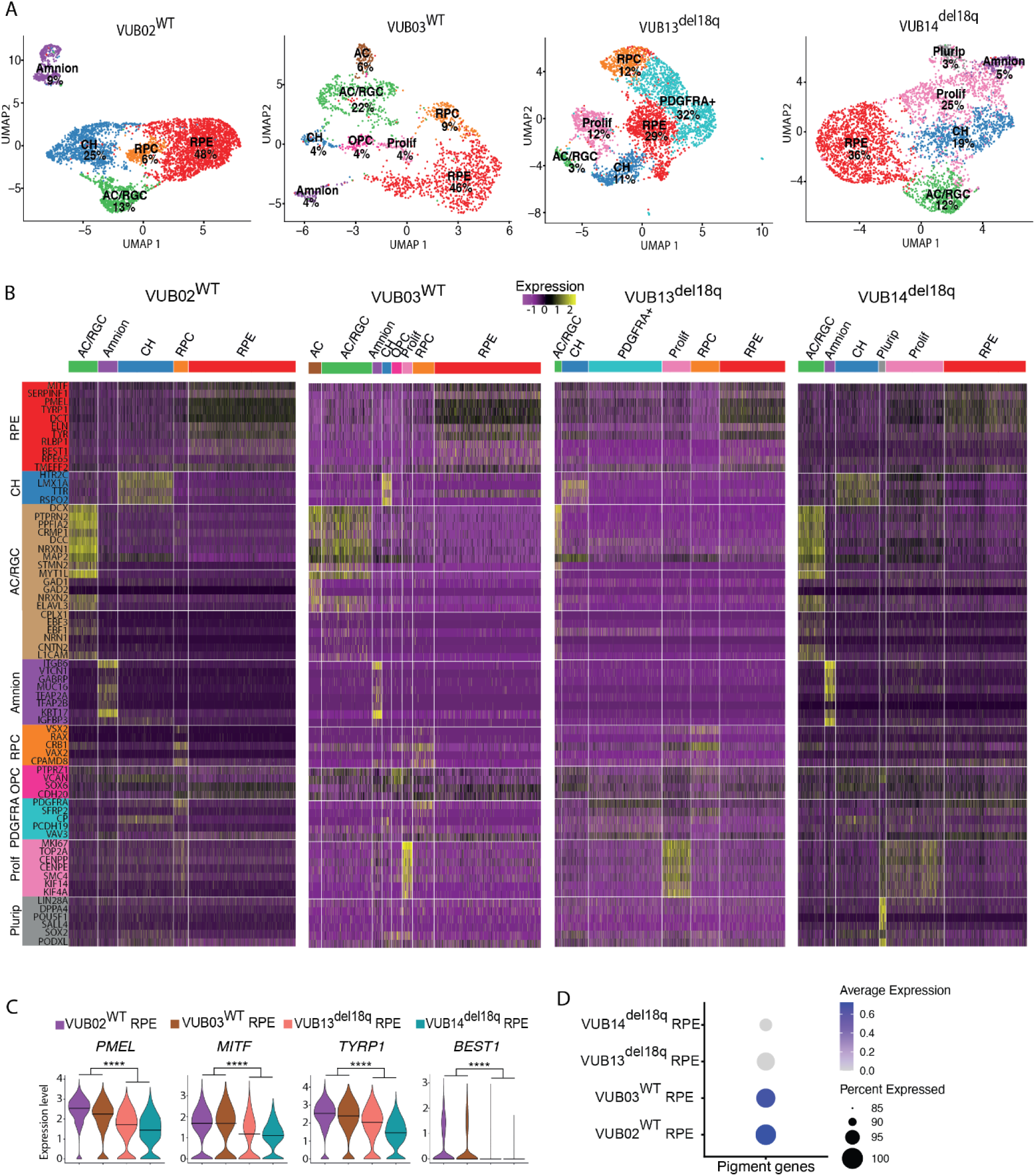
Single-cell RNA sequencing shows that hESCs^del18q^ differentiate into cell populations containing residual undifferentiated cells, proliferating cells and immature RPEs. **A.** UMAP showing the cell composition of every line after 60 days of spontaneous differentiation, with all the major lineages and their relative percentages indicated. Retinal Pigment Epithelium (RPE), Retinal Progenitor cells (RPC), Developing Neuron (Dev-Neu), Cortical Hem (CH), Amnion Cells (Amnion), Amacrine Cells (AC), Retinal Ganglion Cells (RGC), Pluripotent (Plurip), Proliferating (Prolif) **B.** Marker gene expression heatmap for each of the main identified cell types. **C.** Violin plots showing the expression of the key RPE genes *MITF*, *PMEL*, *TYRP1* and *BEST1* in hESCs^del18q^and hESCs^WT^ -derived RPEs. Data are analysis using FindMarkers in Seurat and shown as the “avg_log2FC”. The *, **, *** and **** represent statistical significance between hESCs^WT^ and hESCs^del18q^ at 5%, 1%, 0.1% and 0.01% respectively (“MAST” test function from Seurat). **D.** Dotplots representing the expression averages of pigmentation associated genes in the RPE subpopulations of hESCs^WT^ and hESCs^del18q^.

### hESCs^del18q^ derived cells show deregulation of TGFβ signalling and the absence of expression of RPE master transcriptional regulators

We undertook differential gene expression analysis and Gene set enrichment analysis (GSEA) of the scRNAseq data to infer the molecular mechanisms responsible for the decreased differentiation potential of the variant lines. We first compared the pure RPC and RPE populations separately with undifferentiated hESCs (data from Couvreu et al, submitted, https://doi.org/10.21203/rs.3.rs-5083824/v1, https://www.researchsquare.com/article/rs-5083824/latest), and observed that both RPC and RPE, whether mutated or wild-type, exhibited negative enrichment for gene sets associated with the TGFβ, SMAD2, SMAD3, and Wnt signaling pathways (Fig. 3A and B, lists can be found in the Sup. Table 4 and 5). Next, we compared RPE to retinal progenitor cells and found higher enrichment of these pathways in RPE (Fig. 3C, lists can be found in the Sup. Table 6), indicating that these pathways are activated during the transition from progenitor cells to the RPE state. Together, these data support a specific pattern of TGFβ and Wnt pathway regulation throughout RPE differentiation; in line with a previous study on a two-stage directed RPE differentiation model, where hESCs were initially differentiated by inhibiting TGFβ and Wnt signaling to generate anterior neural ectoderm/eye-field cells, followed by their activation to drive further differentiation and maturation into RPE ^26^.

**Figure 3.**
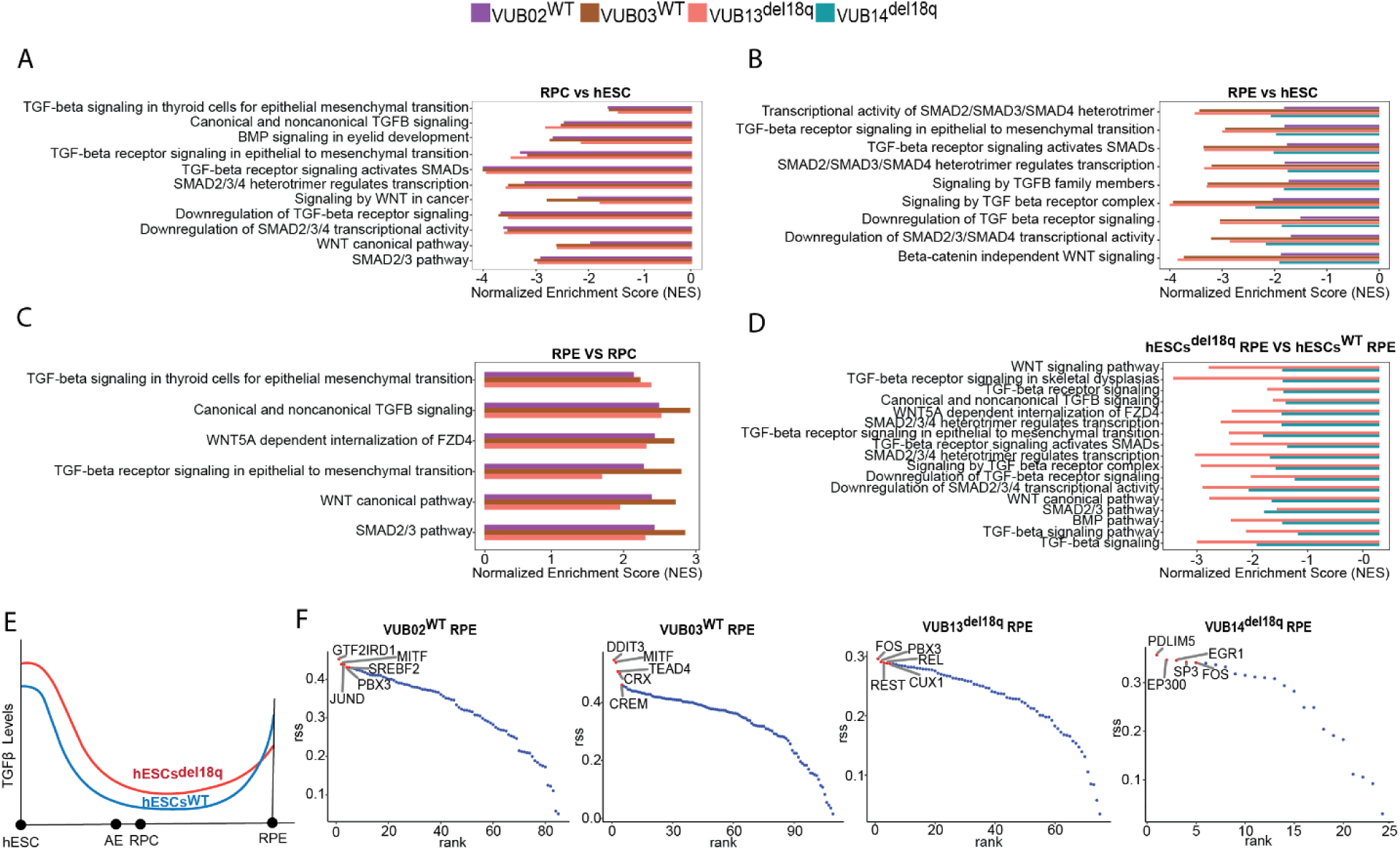
GSEA and SCENIC analysis shows deregulation of the TGFβ and Wnt signaling, and a lack of induction of RPE master regulators in cells with a loss of 18q. **A.** GSEA of the RPC cell population vs hESCs **B.** GSEA of the RPE cell population vs hESCs **C.** GSEA of the RPE vs of RPC **D.** GSEA of the RPE cell population of hESCs^del18q^ vs of hESCs^WT^ **E.** Model of SMADs levels in hESCs^WT^ and hESCs^del18q^ during the whole RPE differentiation **F.** Scenic plot of the master regulators involved in the differentiation to RPE in wt and del18q cells.

We then investigated the transcriptomic differences in the RPE populations from wt and mutant lines that could explain the reduced competence of the mutant cells to correctly and efficiently differentiate into RPE. RPE subpopulations derived from hESCs^del18q^ showed negative enrichment scores for all these TGFβ, SMAD2 and SMAD3 and Wnt signaling pathways (Fig. 3D, lists can be found in the Sup. Table 7), indicating abnormally lower activation of the pathways in hESCs^del18q^ during the maturation from RPC to RPE. Fig. 3E shows a schematic overview of the predicted levels of TGFβ signaling activation at each stage of differentiation in hESCs^WT^and hESCs^del18q^. Last, we employed single-cell regulatory network inference and clustering (SCENIC) analysis to infer gene regulatory network (GRN) sets and the predict transcription factors (TFs) specifically active within each distinct RPE subpopulation. One of the most prominent GRNs in the RPE population is an MITF-dependent network (Fig. 3F). These expected regulons for MITF were active in the hESCs^WT^-RPE, but were notably absent in the hESCs^del18q^-RPE.

### Endogenous overactivation of activin/nodal and Wnt signaling in hESCs**^18q^** leads to abnormal responses in anterior/posterior neuroectoderm differentiation cues

Finally, to unravel the mechanisms behind the differentiation impairment of hESCs^18q^, we investigated the fate of hESCs^WT^ upon spontaneous differentiation and directed neuroectoderm differentiation using Activin/Nodal and BMP4 inhibitors (dual SMAD inhibition), along with retinoic acid. We carried out scRNA sequencing at day 7 of spontaneous differentiation and day 8 of the directed differentiation and found that the two yield different subtypes of neuroectoderm (Fig. 4A). While the cells obtained from the spontaneous differentiation induce higher levels of the anterior neuroectoderm makers *SIX3*, *OTX2*, *RAX*, *LHX2* and *HESX1*, the cells emerging from dual SMAD inhibition express rather posterior neuroectoderm genes (*SOX1*, *SOX3*, *HOXB4*, *HOXA1*, *HOXB1* and *DBX2*) (Fig. 4A). *PAX6* showed to be a pan-neuroectoderm marker, with expression in both lineages. We next sought to assess if modulation of the TGFβ and Wnt signaling (both key in the anterior/posterior neuroectoderm specification) could restore the correct specification of the hESCs^del18q^. hESCs^WT^ and hESCs^del18q^ were treated for 8 days with inhibitors of Activin/Nodal signaling (SB431542, SB), BMP4 signaling (LDN-193189, LDN), and Wnt signaling (XAV-939, XAV), either individually or in combination. Following eight days treatment, a subset of cells was fixed for the assessment of the expression of the eye-field progenitor/anterior neuroectoderm markers SIX3 and the posterior neuroectoderm marker SOX1. The remaining dishes were kept in culture for another four weeks and then imaged to assess for pigmentation as a proxy for successful RPE induction.

**Figure 4.**
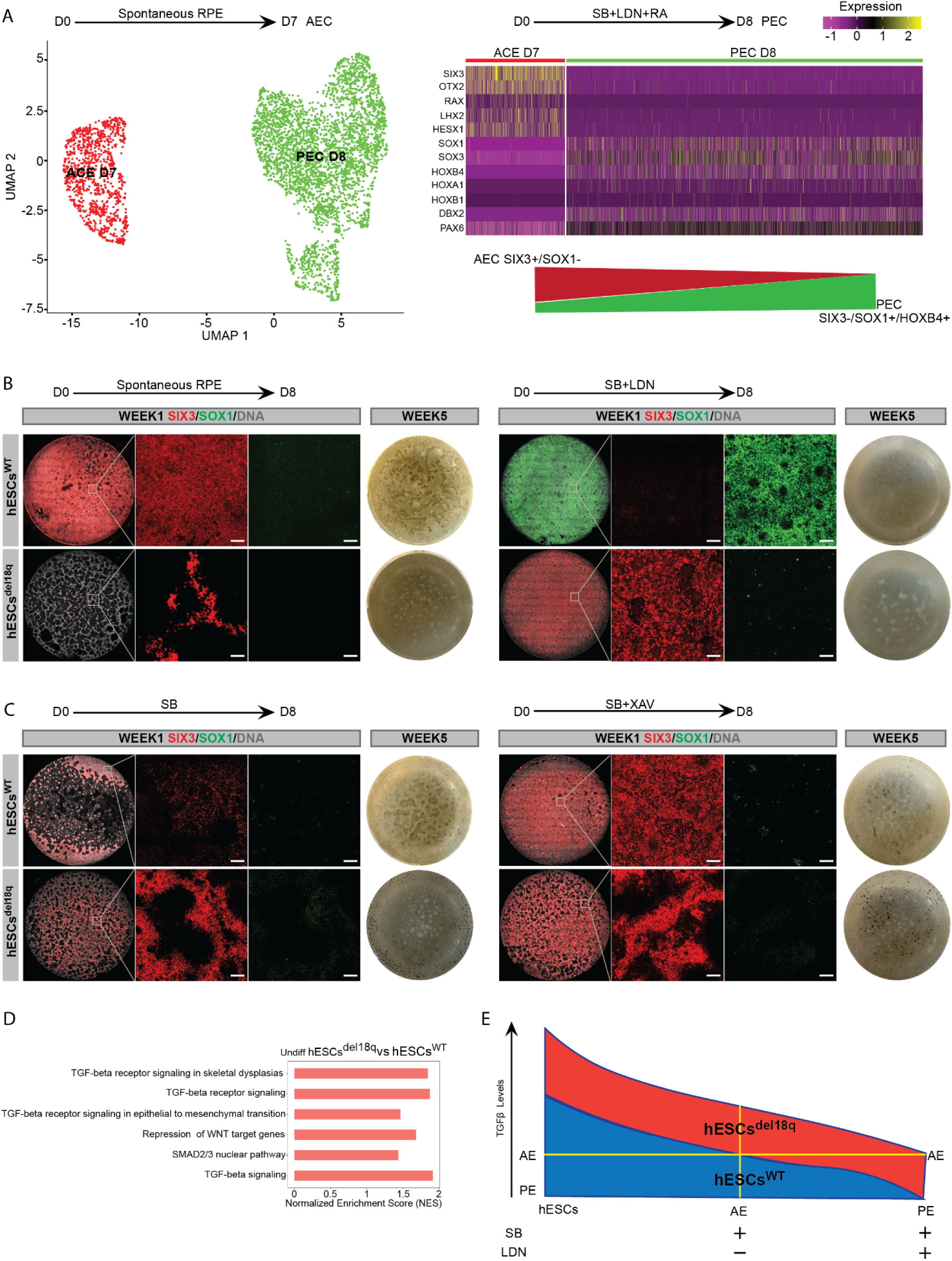
TGFβ and Wnt signaling are at the basis of an improper specification to anterior neuroectoderm and retinal progenitors in cells with a deletion of 18q. **A.** (left) UMAP of the scRNA sequencing of hESCs differentiated to neuroectoderm spontaneously (ACE_D7) or through dual SMAD inhibition and retinoic acid exposure (PEC_D8). (right) The expression of posterior and anterior neuroectoderm markers in the two cell populations reveals two different cell identities.(B-C) Staining of SIX3 (red) and SOX1 (green) on week 1 of spontaneous or dual SMAD inhibition differentiation and whole 1.9 cm^2^ culture plate images to show the RPE pigmentation after 5 weeks differentiation for hESCs^WT^ and hESCs^del18q^. Scale bars of the magnification images are 100 µm (**B**)(left) Spontaneous and (right) dual SMAD inhibition (**C**) (left) SB only (Activin/Nodal inhibition) and (right) SB and XAV (Wnt inhibition). **D.** Normalized enrichment scores for the GSEA for TGFβ and Wnt signalling pathways, comparing the hESCs^del18q^ to hESCs^WT^ in the undifferentiated stage. **E.** Model for the proposed mechanism behind the cell fate differences. Cells in the undifferentiated state have a high activation of the TGFβ signaling. Removal of TGFβ from the culture medium is sufficient to decrease this pathway’s activity to enable anterior neuroectoderm (AE) specification in genetically balanced cells. Further active inhibition of the pathway results in posterior neuroectoderm (PE). Mutant cells have higher endogenous TGFβ activation, and factor withdrawal is insufficient to activate differentiation, and additional pathway inhibition yields only the threshold for anterior neuroectoderm induction.

As seen from the sc-RNAseq, in the hESCs^WT^ spontaneous differentiation without addition of factors resulted predominantly in a *SIX3*-positive *SOX1*-negative anterior neuroectoderm population, while hESCs^del18q^ yielded a poor overall induction to neuroectoderm, with a few *SIX3*-positive cells and no *SOX1*-positive cells (Fig. 4B and Sup. Fig. 2A). Treatment with SB and LDN yielded posterior neuroectoderm in the WT cells, and remarkably, anterior neuroectoderm in the del18q cells. However, these anterior neuroectoderm failed to progress into pigmented RPE at week 5 (Fig. 4B and Sup Fig. 2A). Activin/nodal inhibition through SB treatment alone moderately promoted anterior neuroectoderm induction in del18q cells, which subsequently developed modest black pigmentation by week 5. In contrast, hESCs^WT^ cells exhibited a reduced population of SIX3-positive cells following SB treatment at week 1, along with a notable decrease in pigmentation by week 5 (Fig. 4C and Sup. Fig. 2B). The addition of WNT inhibition with SB did modestly improve anterior neuroectoderm induction in del18q cells, leading to RPE pigmentation by week 5. In comparison, hESCs^WT^ cells demonstrated efficient SIX3 induction by week 1, but exhibited reduced efficiency in RPE induction at week 5 (Fig.4C and Sup. Fig. 2B). Together, this supports the known key role of Activin/Nodal and Wnt signaling in the anterior/posterior axis formation, and suggests that, under normal conditions, BMP4 inhibition is necessary for the formation of posterior neuroectoderm. Conversely, del18q cells require both activin/nodal inhibition to progress to anterior neuroectoderm and Wnt inhibition to generate retinal progenitor cells able to yield correctly pigmented RPE.

Last, in previous work we carried out bulk RNA sequencing of undifferentiated hESCs and found that hESCs^del18q^ display endogenously higher activation of the TGFβ and Wnt signalling as compared to hESCs^WT^(reanalysis shown in Fig. 4D). This is in line with the findings GSEA analysis from the scRNA seq data in this manuscript, where TGFβ and SMAD2/3 playing an important role during the RPE development (Fig. 3E). Taken together, this yields a model in which in WT cells, withdrawal of TGFβ from the culture medium is sufficient to decrease the pathway activation to induce anterior neuroectoderm. Further suppression of the pathway using BMP inhibitors (LDN) results in posterior neuroectoderm fate. In hESCs^18q^, spontaneous differentiation does not result in sufficient suppression of the TGFβ to correctly induce any neuroectoderm specification, and the further inhibitors with LDN only suppress the pathway to meet the threshold for anterior neuroectoderm induction (Fig. 4E).

## Discussion

In this study, we present a systematic evaluation of the impact of segmental losses of chromosome 18 on the developmental trajectory of hESCs differentiating in RPE, using two hESCs^del18q^ lines along with multiple genetically balanced counterparts. Our findings unveil that hESCs harboring an 18q deletion exhibited abnormal specification of the anterior and posterior neuroectoderm due to endogenous hyperactivation of the activin/nodal signaling pathway. This translated into a diminished capacity for RPE differentiation, manifested by sparse pigmented foci and attenuated expression of RPE-specific markers at the endpoint of differentiation. Intriguingly, even in instances where pigmented RPE was formed, these cells exhibited poor RPE characteristics, further highlighting the profound impact of the 18q deletion on both the efficiency and quality of differentiation.

As for the mechanism responsible for this impaired differentiation, we found two key bottlenecks. We observe both an early impairment in the lineage specification of the anterior neuroectoderm, as indicated by the delayed expression of *SIX3*, and an incapacity to fully mature the retinal precursor cells to RPE. This is supported by the results of tracking the developmental process through time-course experiments, the presence of a high proportion of proliferating and pluripotent cells in the scRNAseq, as well as by the significant differences in the transcriptome of the cells. Building upon these insights, we further demonstrated that modulating TGFβ and Wnt pathways through strategic inhibitor treatments could at least partially restore the RPE differentiation of hESCs^del18q^. We found that hESCs^del18q^ respond differently to the TGFβ inhibition than hESCs^WT^. In wild type lines, we found that the balance of specification of anterior and posterior neurectoderm depended on the intensity of the inhibition of the TGFβ signaling. For wild type lines, withdrawal of TGFβ from the culture medium is sufficient to induce anterior neuroectoderm, while additional active inhibition of the two TGFβ pathways drives them to the posterior fate. Further, all treatment conditions involving the BMP inhibitor LDN-193189 blocked RPE maturation in hESCs^WT^ lines, indicating that the additional inhibition of the BMP pathway impedes the normal progression of the RPE differentiation program. In contrast, hESCs^del18q^ lines require additional active inhibition of the pathways to be able to exit the pluripotent state and enter the anterior neuroectoderm lineage. We propose that this is likely due to an endogenously hyperactivated activin/nodal signaling, as already observed in a previous study ^19^

The SCENIC regulon analysis provided insight on what may be a driver of the second barrier to differentiation: the lack of activation of the *MITF* regulon in hESCs^del18q^. *MITF* was shown to be not only a specific marker of the RPE cell population but also to play a crucial functional role as an essential modulator of pigmentation and maturation^27,28^. In line with the findings that Wnt inhibition did enhance the ability of hESCs^del18q^ to generate pigmented RPE, *MITF* has been shown to be regulated by β-catenin and Wnt is known to play a key role in RPE differentiation ^28^. Further research is needed to elucidate the exact nature of the interaction between Wnt signaling and *MITF* in the mutant cells.

Finally, we observed that the differentiated population of hESCs^del18q^ retained a high percentage of proliferating cells, and in one out two lines, we detected also residual pluripotent cells. The presence of these cells would pose an important risk in differentiated products used in transplantation. It is remarkable that a subpopulation of the cells was able to remain in the undifferentiated state 60 days after withdrawal of the factors that are key for pluripotency maintenance and indicates that at least a subgroup of these cells has become growth-factor independent.

There is an increasing body of research focusing on understanding the impact of recurrent genetic abnormalities on the characteristics and function of hPSCs. These abnormalities can undermine their quality and viability for future applications by altering their growth patterns and differentiation potential. For instance, gain of 20q11.21 has been extensively studied, with substantial evidence indicating that hPSCs harboring this alteration altered the distribution of lineages differentiation in embryoid bodies and hinder the commitment to the neuroectodermal lineage^29,30^. A recent study demonstrated that hPSCs with an isochromosome 20q (iso20q) are prone to apoptosis and are unable to differentiate into the primitive germ layers during spontaneous RPE differentiation. In this study we provide novel mechanistic insight into the impact of aneuploidy on hESCs differentiation in a clinically relevant cell type and show that differential activity of TGFβ is at the basis of the anterior/posterior neuroectoderm specification in hESCs. Our work contributes to the mapping of the risks associated to genetic abnormalities in hPSC, and highlights the importance of thorough genetic testing of cells prior to their use in research and in a clinical setting.

## Data availability

Raw sequencing data of human samples is considered personal data by the General Data Protection Regulation of the European Union (Regulation (EU) 2016/679), because SNPs can be extracted from the reads, and cannot be publicly shared. The data can be obtained from the corresponding author upon reasonable request and after signing a Data Use Agreement. sc-RNA sequencing data supporting the figures in this paper can be found at the Open Science Framework, as well as all the data supporting all figures in this paper (https://osf.io/cnvgt/).

## Resource and Materials availability

Further information and requests for resources should be directed to the corresponding author, Claudia Spits.

All VUB stem cell lines in this study, including the genetically abnormal sublines and genetically modified lines, are available upon request and after signing a material transfer agreement.

## Ethics statement

For all parts of this study, the design and conduct complied with all relevant regulations regarding the use of human materials, and all were approved by the local ethical committee of the University Hospital UZ Brussel and the Vrije Universiteit Brussel (File number: B.U.N. 1432020000284). All patients donating embryos to derive human embryonic stem cell lines gave written consent.

## Acknowledgments

Y.L. is a predoctoral fellow supported by the China Scholarship Council (CSC), and M.R., C.J., N.K. and E.C.D.D. are predoctoral fellows supported by the Fonds voor Wetenschappelijk Onderzoek Vlaanderen (FWO). M.C.D. is a predoctoral fellow supported by the 175 Military Hospital in Vietnam. This research was supported by the FWO (grant numbers 1506617N and G0713222N) and the Methusalem Grant to Karen Sermon and Claudia Spits (Vrije Universiteit Brussel). The authors wish to thank An Verloes and Brecht Ghesquiere for their support to this study.

## Author contributions

Y.L. carried out all the experiments and bioinformatics analysis unless stated otherwise and co-wrote the manuscript. C.M.D. and A.H assisted with the mRNA extraction and qPCR. D.A.D, C.J., N.K. and C.M.D. assisted with cell culture and immunostaining, E.C.D.D. assisted with the bioinformatics analysis. M.R. assisted in microscopy and making of the figures. K.S. proofread the paper. C.S. co-wrote the manuscript and designed and supervised the experimental work.

## References

1 Zarbin M. Cell-based therapy for degenerative retinal disease. Trends Mol Med. 2016; 22: 115– 134.

2 da Cruz L, Chen FK, Ahmado A, Greenwood J, Coffey P. RPE transplantation and its role in retinal disease. Prog Retin Eye Res 2007; 26: 598–635.

3 Mehat MS, Sundaram V, Ripamonti C, Robson AG, Smith AJ, Borooah S et al. Transplantation of Human Embryonic Stem Cell-Derived Retinal Pigment Epithelial Cells in Macular Degeneration. Ophthalmology 2018; 125: 1765–1775.

4 Schwartz SD, Regillo CD, Lam BL, Eliott D, Rosenfeld PJ, Gregori NZ et al. Human embryonic stem cell-derived retinal pigment epithelium in patients with age-related macular degeneration and Stargardt’s macular dystrophy: Follow-up of two open-label phase 1/2 studies. The Lancet 2015; 385: 509–516.

5 Da Cruz L, Fynes K, Georgiadis O, Kerby J, Luo YH, Ahmado A et al. Phase 1 clinical study of an embryonic stem cell-derived retinal pigment epithelium patch in age-related macular degeneration. Nat Biotechnol 2018; 36: 328–337.

6 O’Neill HC, Limnios IJ, Barnett NL. Advancing a Stem Cell Therapy for Age-Related Macular Degeneration. Curr Stem Cell Res Ther 2019; 15: 89–97.

7 Schwartz SD, Regillo CD, Lam BL, Eliott D, Rosenfeld PJ, Gregori NZ et al. Human embryonic stem cell-derived retinal pigment epithelium in patients with age-related macular degeneration and Stargardt’s macular dystrophy: follow-up of two open-label phase 1/2 studies. The Lancet 2015; 385: 509–516.

8 Vugler A, Carr AJ, Lawrence J, Chen LL, Burrell K, Wright A et al. Elucidating the phenomenon of HESC-derived RPE: Anatomy of cell genesis, expansion and retinal transplantation. Exp Neurol 2008; 214: 347–361.

9 Andrews PW, Barbaric I, Benvenisty N, Draper JS, Ludwig T, Merkle FT et al. The consequences of recurrent genetic and epigenetic variants in human pluripotent stem cells. Cell Stem Cell 2022; 29: 1624–1636.

10 Amps K, Andrews PW, Anyfantis G, Armstrong L, Avery S, Baharvand H et al. Screening ethnically diverse human embryonic stem cells identifies a chromosome 20 minimal amplicon conferring growth advantage. Nat Biotechnol 2011; 29: 1132–1144.

11 Merkle FT, Ghosh S, Kamitaki N, Mitchell J, Avior Y, Mello C et al. Human pluripotent stem cells recurrently acquire and expand dominant negative P53 mutations. Nature 2017; 545: 229–233.

12 Martins-Taylor K, Nisler BS, Taapken SM, Compton T, Crandall L, Montgomery KD et al. Recurrent copy number variations in human induced pluripotent stem cells. Nat Biotechnol 2011; 29: 488– 491.

13 Taapken SM, Nisler BS, Newton MA, Sampsell-Barron TL, Leonhard KA, McIntire EM et al. Karotypic abnormalities in human induced pluripotent stem cells and embryonic stem cells. Nat Biotechnol. 2011; 29: 313–314.

14 Merkle FT, Ghosh S, Genovese G, Handsaker RE, Kashin S, Meyer D et al. Whole-genome analysis of human embryonic stem cells enables rational line selection based on genetic variation. Cell Stem Cell 2022; 29: 472–486.e7.

15 Keller A, Spits C. The Impact of Acquired Genetic Abnormalities on the Clinical Translation of Human Pluripotent Stem Cells. Cells 2021; 10. doi:10.3390/CELLS10113246.

16 Keller A, Dziedzicka D, Zambelli F, Markouli C, Sermon K, Spits C et al. Genetic and epigenetic factors which modulate differentiation propensity in human pluripotent stem cells. Hum Reprod Update 2018; 24: 162–175.

17 Halliwell J, Barbaric I, Andrews PW. Acquired genetic changes in human pluripotent stem cells: origins and consequences. Nat Rev Mol Cell Biol. 2020; 21: 715–728.

18 Al Delbany D, Ghosh MS, Krivec N, Huyghebaert A, Regin M, Duong MC et al. De Novo Cancer Mutations Frequently Associate with Recurrent Chromosomal Abnormalities during Long-Term Human Pluripotent Stem Cell Culture. Cells 2024; 13: 1395.

19 Lei Y, Al Delbany D, Krivec N, Regin M, Couvreu de Deckersberg E, Janssens C et al. SALL3 mediates the loss of neuroectodermal differentiation potential in human embryonic stem cells with chromosome 18q loss. Stem Cell Reports 2024; 19: 562–578.

20 Mateizel I, Spits C, De Rycke M, Liebaers I, Sermon K. Derivation, culture, and characterization of VUB hESC lines. In Vitro Cell Dev Biol Anim 2010; 46: 300–308.

21 Mateizel I, De Temmerman N, Ullmann U, Cauffman G, Sermon K, Van de Velde H et al. Derivation of human embryonic stem cell lines from embryos obtained after IVF and after PGD for monogenic disorders. Hum Reprod 2006; 21: 503–511.

22 Plaza Reyes A, Petrus-Reurer S, Padrell Sánchez S, Kumar P, Douagi I, Bartuma H et al. Identification of cell surface markers and establishment of monolayer differentiation to retinal pigment epithelial cells. Nat Commun 2020; 11. doi:10.1038/s41467-020-15326-5.

23 Plaza A, Sandra R, Sara P-R, Sánchez P, Kumar P, Douagi I et al. Xeno-free, chemically dened and scalable monolayer differentiation protocol for retinal pigment epithelial cells. 2020. doi:10.21203/rs.3.pex-635/v1.

24 Plaza Reyes A, Petrus-Reurer S, Antonsson L, Stenfelt S, Bartuma H, Panula S et al. Xeno-Free and Defined Human Embryonic Stem Cell-Derived Retinal Pigment Epithelial Cells Functionally Integrate in a Large-Eyed Preclinical Model. Stem Cell Reports 2016; 6: 9–17.

25 Bayindir B, Dehaspe L, Brison N, Brady P, Ardui S, Kammoun M et al. Noninvasive prenatal testing using a novel analysis pipeline to screen for all autosomal fetal aneuploidies improves pregnancy management. Eur J Hum Genet 2015; 23: 1286–1293.

26 Limnios IJ, Chau Y-Q, Skabo SJ, Surrao DC, O’Neill HC. Efficient differentiation of human embryonic stem cells to retinal pigment epithelium under defined conditions. Stem Cell Res Ther 2021; 12: 248.

27 Capowski EE, Simonett JM, Clark EM, Wright LS, Howden SE, Wallace KA et al. Loss of MITF expression during human embryonic stem cell differentiation disrupts retinal pigment epithelium development and optic vesicle cell proliferation. Hum Mol Genet 2014; 23: 6332–6344.

28 Westenskow P, Piccolo S, Fuhrmann S. β-catenin controls differentiation of the retinal pigment epithelium in the mouse optic cup by regulating Mitf and Otx2 expression. Development 2009; 136: 2505–2510.

29 Jo HY, Lee Y, Ahn H, Han HJ, Kwon A, Kim BY et al. Functional in vivo and in vitro effects of 20q11.21 genetic aberrations on hPSC differentiation. Sci Rep 2020; 10: 1–15.

30 Markouli C, De Deckersberg EC, Regin M, Nguyen HTT, Zambelli F, Keller A et al. Gain of 20q11.21 in Human Pluripotent Stem Cells Impairs TGF-β-Dependent Neuroectodermal Commitment. Stem Cell Reports 2019; 13: 163–176.

