## Supplementary data for "Loss of 18q alters TGFβ signalling affecting anteroposterior neuroectodermal fate in human embryonic stem cells"

**Supplemental Data**

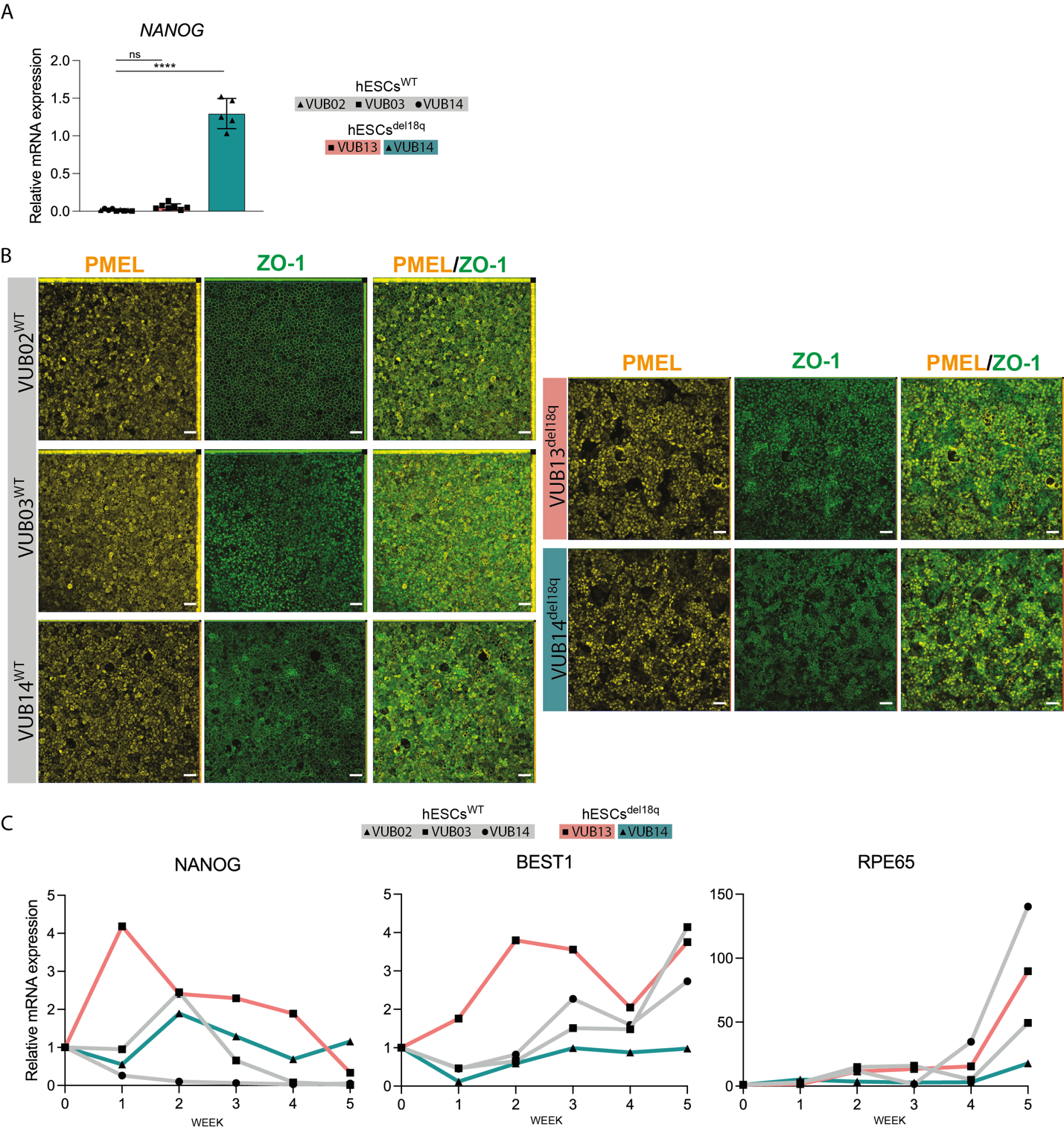

**Supplemental Figure 1**. **hESCs with a loss of 18q show a reduced differentiation potential to RPE and an overall poor anterior neuroectoderm induction. A**. Relative mRNA expression of pluripotency (*NANOG*) markers of the dishes shown in A, prior to RPE purification. Data information: Data are shown as the means ± SEM (hESCs^WT^: N=3/per line, VUB13^del18q^: N = 6, VUB14^del18q^: N = 5). Each datapoint refers to an independent differentiation experiment and *, **, *** and **** represent statistical significance between samples at 5%, 1%, 0.1% and 0.01% respectively (unpaired t-test).**B.** Staining of purified RPE for the mature RPE marker PMEL and ZO-1, all scale bars are 50 µm. **C**. Dynamics of the relative mRNA expression of *NANOG*, *BEST1 and RPE65* at 5 time points (W1, W2, W3, W4, W5) during RPE differentiation.

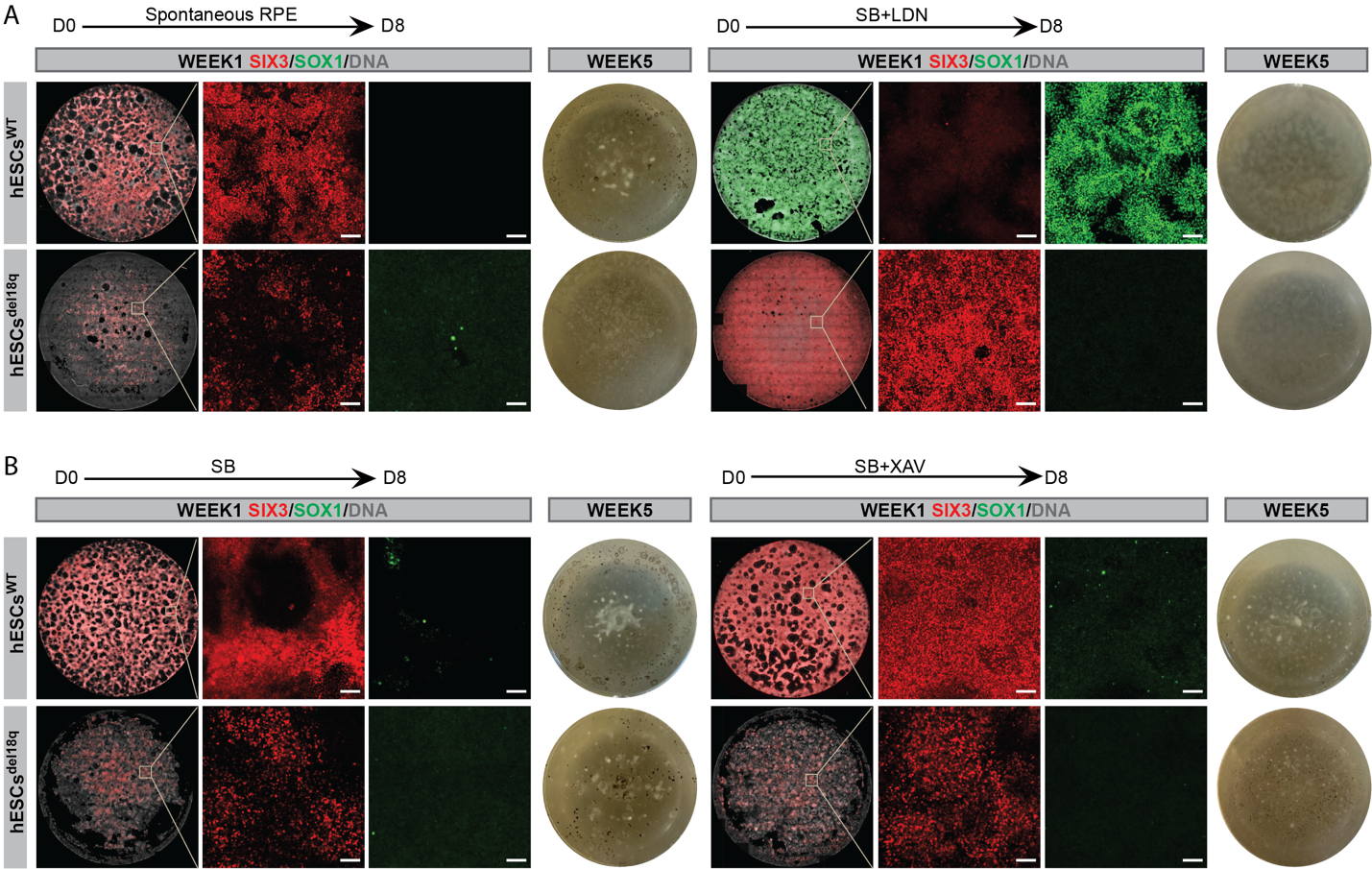

**Supplemental Figure 2. TGFβ and Wnt signaling are at the basis of an improper specification to anterior neuroectoderm and retinal progenitors in cells with a deletion of 18q. A-B.** Staining of SIX3 (red) and SOX1 (green) on week 1 of spontaneous or dual SMAD inhibition differentiation and whole 1.9 cm^2^ culture plate images to show the RPE pigmentation after 5 weeks differentiation for hESCs^WT^ and hESCs^del18q^. Scale bars of the magnification images are 100 µm (**B**)(left) Spontaneous and (right) dual SMAD inhibition (**C**) (left) SB only (Activin/Nodal inhibition) and (right) SB and XAV (Wnt inhibition).

**Supplemental Table 1. List of genes used to identify the related cell clusters for figure 2B and dotplot for Figure2D**

| **Cell type** | **Genes** |
| --- | --- |
| RPE | MITF/SERPINF1/PMEL/TYRP1/DCT/ELN/TYR/RLBP1/BEST1/RPE65/TMEFF2 |
| hem | HTR2C/LMX1A/TTR/RSPO2/PCP4/CLDN5 |
| Dev_Neu | DCX/PTPRN2/PPFIA2/CRMP1/DCC/NRXN1/MAP2/STMN2 |
| Amacrine | MYT1L/GAD1/GAD2/NRXN2/ELAVL3 |
| hem1 | CPLX1/EBF3/EBF1/NRN1/CNTN2/L1CAM |
| Amnion | ITGB6/VTCN1/GABRP/MUC16/TFAP2A/TFAP2B/KRT17/IGFBP3 |
| RPC | VSX2/RAX/CRB1/VAX2/CPAMD8 |
| oligodendrocyte_progenitor | PTPRZ1/VCAN/SOX6/CDH20 |
| proliferating | MKI67/TOP2A/CENPF/CENPE/SMC4/KIF14/KIF4A |
| pluripotent | LIN28A/DPPA4/POU5F1/SALL4/SOX2/PODXL |
| extraembryonic_mesoderm | LUM/NID2/FOXF1/VIM/POSTN/ANXA1/PITX1 |
| Mesenchyme | COL3A1/PITX1/LUM/HAND1 |
| Schwann_cell | FOXD3/TFAP2B/SOX10/ERBB3 |
| Neuroblasts | VAX1/ENTPD1/BRINP2/ANGPTL1/AMBN/EMX1/NEUROG2/POU3F3 |
| Sensory_Neurons | ATOH1/OLIG3/PDE1A/POU3F4 |
| Muller_glia | PLP1/TF/RLBP1/FABP7/WIF1/LGI4/SLC1A3/SPON1/PMEPA1 |
| Microglia | CX3CR1/C1QA/C1QB/C1QC/TYROBP/AIF1/LAPTM5/ITGB2/RNASET2/  CSF2RA/FCER1G/S100A11/CSF1R |
| bipolar_cell | VSX1/VSX2/RPDM8/GRM6/NEUROD4/CA10/PLXDC1 |
| horizontal_cell | ONECUT1/ONECUT2/ONECUT3/SULF2/PROX1 |
| photoreceptor | PDC/PDE6G/SAG/CRX/NRL/RCVRN/AIPL1/PDC/NRL/ROM1/NR2E2/RP1 |
| Astrocyte | AQP4/SLC1A3/GLUL/GFAP |
| Pericytes | ACTA2/TPM2/C11orf96/HIGD1B/TAGLN/RGS5 |
| Cone | MYL4/GUCA1C/OPN1LW/ARR3/PDE6H |
| Rod | REEP6/DRAIC/CLUL1/NR2E3/PDE6A/CNGA1 |
| Pigmentation | TYR/DCT/TYRP1/SLC45A2/MITF/OCA2/GPR143/SLC24A5/KIT/MC1R/KLF6  /COMMD3 |

**Supplementary Table 2. Taqman assays (Thermofisher) used in RT-qPCR**

| **Gene** | **Reference** |
| --- | --- |
| *BEST1* | Hs00188249_m1 |
| *RPE65* | Hs00165642_m1 |
| *PMEL* | Hs00173854_m1 |
| *MITF* | Hs01117294_m1 |
| *PAX6* | Hs00240871_m1 |
| *SIX3* | Hs00193667_m1 |
| *RAX* | Hs00429459_m1 |
| *POU5F1* | Hs00742896_s1 |
| *NANOG* | Hs02387400_g1 |
| *ID1 – copy number* | Hs01892845_cn |
| *KIF14– copy number* | Hs00637799_cn |
| *NANOG– copy number* | Hs03820140_cn |
| *RNASEP – copy number* | Hs4403328_cn |

**Supplementary Table 3: Primary and secondary antibodies**

| **Primary antibodies** | **Reference** |
| --- | --- |
| BEST1 | Merck Millipore, MAB5466 |
| PAX6 | Invitrogen, SD08-31 |
| PMEL | Invitrogen, MA1-34759 |
| ZO-1 | Invitrogen, 61-7300 |
| SIX3 | Abcam, ab221750 |
| SOX1 | R&D Systems, AF3369-SP |
| **Secondary antibodies** |  |
| Alexa Fluor 488 donkey anti-rabbit IgG (H+L) | Thermofisher, A-21206 |
| Alexa Fluor 647 donkey anti-mouse IgG (H+L) | Thermofisher, A-31571 |
| Alexa Fluor 488 donkey anti-goat IgG (H+L) | Thermofisher, A-11055 |
| Alexa Fluor 594 donkey anti-rabbit IgG (H+L) | Thermofisher, A-21207 |
